# A stimuli-responsive *ex vivo* model of osteoarthritis demonstrates TLR4-mediated cartilage degradation and a Rapamycin-induced fast matrix recovery

**DOI:** 10.64898/2026.06.09.731049

**Authors:** R. Di Gesù, H Kenawy, G. Vitale, I. Chiesa, R. Gottardi

## Abstract

**Background:** In osteoarthritis (OA) TLR4 signaling leads to downstream activation of the phosphoinositide 3-kinases/ protein kinase B/ mammalian target of Rapamycin (PIK3/AKT/mTOR) pathway, a known modulator of autophagic mechanisms in chondrocytes. This paper focuses on creating a realistic *ex vivo* OA model that mimics elements of the pathophysiology of OA, allowing for further hypotheses-based investigations, and for use as a bench test for new therapeutic targets.

**Objective:** To study the downstream inflammatory and matrix changes in cartilage due to TLR4 signaling and the recovery achieved by a commonly used immunosuppressive drug, Rapamycin.

**Methods:** In an *ex vivo* 3D model based on healthy porcine cartilage explants, we mimicked the OA environment by LPS stimulation activating TLR4 signaling. Furthermore, we inhibited mTOR signaling via Rapamycin, which is accepted to attenuate the cartilage response to LPS-TLR4 activation. Histology and immunohistochemistry were used to evaluate the structural and biomolecular modifications driven by LPS and Rapamycin.

**Results:** The explant model captured key features of OA, such as extracellular matrix degeneration and altered autophagy. The OA-like changes in the model were driven by TLR4 activation and mTOR signaling, well-known OA-related molecular pathways, and reversed by Rapamycin.

**Conclusion:** We demonstrate that our explant model is responsive to LPS stimulation, leading to activation of OA-related biomolecular pathways, closely mimicking the native physiological processes. This evidence supports the potential of our model to act as a platform for OA studies, in particular related to the gut-joint axis in age-related OA, and for the screening of new disease-modifying molecules.

## I. Introduction

Osteoarthritis (OA) is a chronic synovial joint disorder, causing pain, swelling, and anatomic deformations, ultimately compromising joint function^1^, affecting over 250 million people worldwide^2^, with the elderly population contributing to a higher proportion^3,4^. OA correlates with aging and comorbidities by an incidence of up to 20%^5,6^. This pathology leads to invalidating conditions which seriously compromises daily activities resulting in management through supportive assistance^7^. Most treatments focus on pain controlling via NSAIDs and corticosteroid injections until total joint replacement is necessary^8^. The most obvious manifestation of OA is articular cartilage damage which is often associated with extracellular matrix (ECM) dysplasia through a decreased type II / type I collagen ratio^9^ and an increase in glycosaminoglycan (GAG) loss^10^. However, the heterogenous mechanisms causing OA have yet to be fully understood.

One mechanism that has been implicated in the progression of OA is Toll-like receptor 4 (TLR4) signaling (**Fig. 1**) supported by co-receptor MD2, that contributes to triggering inflammation potentially via downstream nuclear factor-kappa B (NFkB)^11,12^. NFkB activation has been shown to induce cartilage degradation increasing expression of matrix-metalloproteinases and chondrocyte hypertrophy^13–15^. TLR4 signaling in OA can be activated by multiple ECM-derived DAMPs (e.g., low-molecular-weight hyaluronan fragments, tenascin-C, fibronectin, etc.) whose production is increased in OA in both synovial fluid and joint tissues.TLR4 signaling can also be triggered by lipopolysaccharide (LPS), a gram-negative bacteria-derived endotoxin that can serve as a ligand to TLR4^16^. Due the simplicity of the direct cause and effect relationship between LPS and TLR4, LPS is often used to stimulate inflammation and initiate cartilage matrix degradation^11,12,17^. Furthermore, leveraging LPS to model the effect of TLR4 in cartilage degeneration allows to dissect the response to TLR4 activation orthogonally to the other effects DAMPs may have. Notably, although LPS might not be a TLR4 ligand commonly considered in OA conditions, an association of LPS with OA has been proposed via the leaky gut theory^18^, an age-related OA comorbidity^19–22^. Leaky gut is thought to be caused by an imbalance of the gut-resident microbiota, resulting in a reduction of the beneficial microorganisms subpopulation (dysbiosis) in the gut epithelium^23^. As a consequence, this condition results in the partial or total laxity of enterocytes’ tight junctions^24–26^ which is the cell-cell connection that blocks the paracellular translocation of substances^27^ in the gut (**Fig. S1**). Thus, LPS may be more implicated in osteoarthritic cartilage degeneration than previously believed.

**Figure 1:**
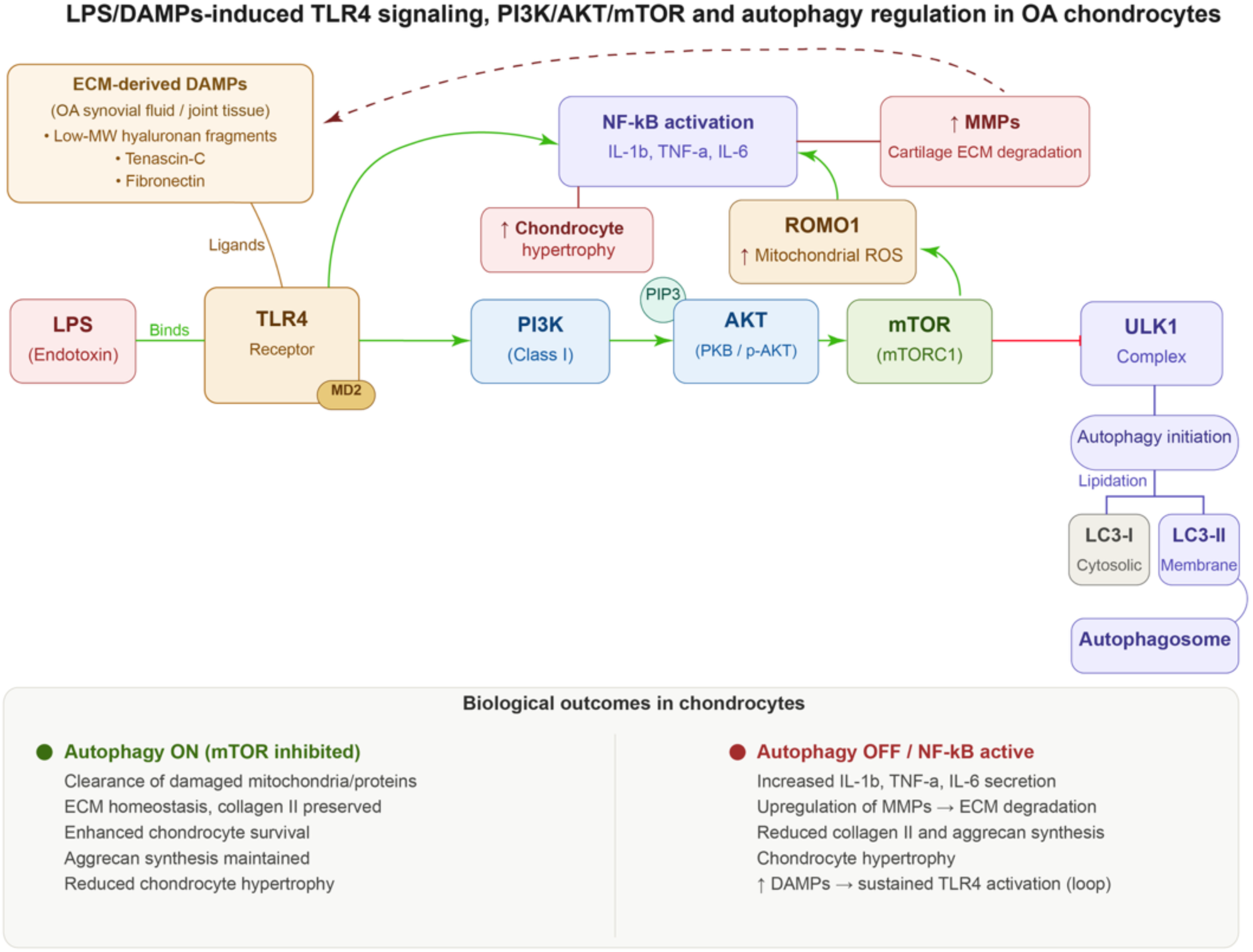
LPS-induced PI3K/AKT/mTOR signaling regulates autophagy in chondrocytes. LPS activates TLR4-PI3K/AKT/mTOR signaling, inhibiting ULK1-mediated autophagy and promoting inflammation; stress-induced mTOR inhibition activates protective autophagy.

Interestingly, TLR4 also activates downstream the phosphoinositide 3-kinases/protein kinase B/mammalian target of Rapamycin (PIK3/AKT/mTOR), modulating autophagy in chondrocytes^28–31^. Autophagy normally regulates cartilage homeostasis by processing damaged cellular components through autophagic vesicles by fusing with lysosomes for end-stage degradation (**Fig. S2**)^32–36^. Many factors often associated with age-related OA may affect the autophagy mechanism such as SIRT1, FOXO3, NFkB, p53, and mTOR. Two protein complexes, mTORC1 and mTORC2 are part of mTOR signaling (**Fig. S3**) and when activated, this pathway leads to inhibition of autophagy. Hence, the LPS-TLR4-mTOR axis^37^ acts as an autophagy repressor, as demonstrated by several findings showing decreased autophagic tendencies of LPS-treated cells and tissues^32,38,39^. LC3b and ULK1 are proteins that strongly sustain autophagy *in vivo*^32,38,40–42^, with ULK1 playing a key upstream role in the transduction of pro-autophagic signals promoting the autophagosome formation through LC3^43^. Both ULK1 and LC3 are expressed in healthy murine articular cartilage, whereas chondrocytes from surgically-induced OA present decreased ULK1 and LC3 expression^38^, suggesting the need to investigate autophagy in an inflamed cartilage model. Besides systemic inflammation and autophagy, LPS also triggers the production of reactive oxygen species (ROS), a key element of age-related OA. ROS induce inflammatory matrix degradation, calcification, apoptosis, and senescence^44–47^, mainly due to the activation of ROS modulator 1 (ROMO), as well as decrease in cell proliferation^29,48^. Furthermore, ROS can also activate the PIK3/AKT/mTOR, leading to an inflammatory feedback loop^47,48^ (**Fig. S4**).

While there are extensive studies that examine OA-related pathways to identify potential drug targets, no resolutive pharmacologic therapy has yet been identified. Rapamycin, an antifungal antibiotic^49^, is a commonly used immunosuppressive drug that serves as a mTOR inhibitor, leading to the restoration of native cellular autophagy. Rapamycin has been shown to restore autophagy in DMM-induced OA *in vivo*^50,51^, as well as in human-derived chondrocytes where it attenuated OA and prevented senescence^52^. Furthermore, there are also a few studies that examine the effect of Rapamycin on OA-related inflammation both *ex vivo*^53^ an *in vivo* in mouse models^54^. However, although Rapamycin has been used in preclinical models, there are no clinical trials studying its effect on OA cartilage.

Given the ECM-rich nature of cartilage, 2D *in vitro* models of OA have had limited success in recapitulating the *in vivo* environment to study cartilage degradation^55–59^, with few exceptions^60^. *Ex vivo* explants as we have implemented in this work, represent then an excellent alternative to model OA *in vitro* capturing the 3D and ECM-rich nature of cartilage^55–59^. Furthermore, to the best of our knowledge, *ex vivo* OA models when not derived from mechanical overloading^1^, have so far relied on direct stimulations by pro-inflammatory cytokines such as IL-1β or TNF-α^53^, which induce autocrine feedback loops making it arduous to distinguish the effects of the exogenous cytokines versus the autocrine inflammatory response.

To overcome these limitations, in our study, we mimic the OA environment using an *ex vivo* model based on healthy porcine cartilaginous explants stimulated with LPS. Thus, via LPS stimulation, we drive and OA-like state via TLR4 activation. Furthermore, it is expected that we would be able to inhibit mTOR signaling via Rapamycin, attenuating the cartilage response to LPS-TLR4 activation.

**Our objectives were to study the downstream inflammatory and matrix changes in cartilage due to TLR4 signaling and compare the morphological changes in cartilage post LPS insult as well as in recovery after treatment with Rapamycin.** Through this approach, we aim to overcome limitations of routinely adopted protocols based on stimulation with pro-inflammatory cytokines^61,62^, and rather, adopt LPS as the single inflammatory-trigger to induce an OA-like state via TLR-4 pathway activation^11^. Thus, our work aims to create a realistic 3D OA model that mimics the pathophysiology of OA, allowing for further hypothesis-driven investigations, and for the future screening of new therapeutic targets.

## II. Methods

### Cartilage harvest

Cartilage plugs were harvested from intact juvenile porcine (*sus scrofa,* 3-6 months old) knees obtained from a local abattoir within 24 hours *post mortem*. Cartilage plugs were harvested from femoral condyles using disposable biopsy punches (Ø = 4mm, Kai medical, Japan) maintaining a cartilage surface-punch angle of 90° to obtain cylinder-shaped discs.

### Cartilage explant culture

Plugs (N=3 biological donors) were cultured in nontreated 24 well plates at 37 °C and 5% CO_2_ in chondrogenic medium (CMM: Fluorobrite-Gibco, USA, L-Proline 40 µg/ml-Sigma, USA, GlutaMax 1% v/v-Gibco, USA, sodium pyruvate 1% v/v-Sigma, USA, antibiotic-antimycotic solution 2% v/v-Gibco, USA, insulin-transferrin-selenium 1% v/v-Sigma, USA, TGF-β3 10 ng/ml-PeproTech, UK, ascorbic acid 50 µg/ml-Sigma, USA)) for 24h to allow for recovery post-harvest. At day 0 (D0), plugs were divided in 3 groups and cultured as follows: the **control group (-L-R)** was cultured for 7 days in CMM, the LPS **(inflamed) group (+L-R)** was cultured in CMM containing LPS (Sigma, USA) (CMM-LPS) for 3 days (D3), then switched to CMM up to (D7), and the **LPS+treatment group (+L+R)** was cultured with CMM-LPS up to D3, and then switched to CMM containing Rapamycin^17^ (Sigma, USA) (CMM-R) up to D7. For each group, 2 plugs were collected at each time point (D0 and D7) and condition for histological preparation.

### Paraffin embedding and sectioning

Plugs (N=18, 3 donors for 3 conditions for 2 time points) were fixed overnight with 4% paraformaldehyde solution (Sigma, USA) at 4°C under constant mechanical agitation. After dehydration in ascending EtOH series (25% - 100%) followed by xylene (Sigma, USA), samples were embedded in histology-grade paraffin (Leica biosystems, Germany) using the HistoCore Arcadia H embedding station (Leica biosystems, Germany). Embedded samples were sectioned at 5 μm thickness using a fully-motorized rotary microtome Leica RM2265 (Leica biosystems, Germany), collected on superfrost plus adhesion slides (Thermofisher, USA), and stored at room temperature (RT). For histochemical and immunohistochemical staining, the sections were preliminarily de-paraffinized with xylene, rehydrated in descending EtOH series (100% - 50%), and gently washed under running cold tap water.

### Histochemistry

For Hematoxylin and Eosin (H&E) staining, slides were soaked in a Harris modified hematoxylin solution 2% w/v (Fisher Scientific, USA) for 5 minutes, gently washed under running tap water for 2 minutes, soaked in a 1% w/v Eosin solution (Fisher Scientific, USA) in EtOH 80% v/v - acetic acid 1.25% v/v for 40 seconds. Then, slides were gently washed under running tap water, dehydrated in ascending EtOH series (25%, 50%, 70%, 95%, and 100%), mounted with limonene mounting medium (Sigma, USA), and stored at RT away from light. For Alcian Blue (AB) staining, slides were soaked in an Alcian blue solution 1% w/v in acetic acid (pH = 2.5) (Sigma, USA) for 1 hr at room temperature, and the staining solution was washed away under a gentle flow of running tap water. Then, slides were dehydrated in ascending ethanol series (25%, 50%, 70%, 95%, and 100%), mounted with limonene mounting medium, and stored at RT away from light. For Safranin O/Fast Green (SafO-FG) staining, slides were immersed in a Fast Green (Sigma, USA) 0.1% w/v solution in distilled H_2_0 for 7 minutes at RT, rinsed with 1% v/v acetic acid (Sigma, USA), and stained with a Safranin O (Sigma, USA) 0.1 % w/v solution in distilled H_2_0 for 30 minutes at RT. Hence, slides were dehydrated in ascending EtOH series (25%, 50%, 70%, 95%, and 100%), mounted with limonene mount medium, and stored at room temperature away from light.

### Immunohistochemistry

For immunostaining, Ms-ROMO1 (dilution 1:150, ab236409, Abcam, USA) Rb-ULK1 (dilution 1:50, ab203207, Abcam, USA) Rb-COLII (dilution 1:50, ab34712, Abcam, USA), Ms-LC3β (dilution 1:100, NB100-2220, Novusbio, USA) were used as primary unconjugated antibodies. Donkey anti-Mouse DL550 (dilution 1:50, Ab98795, Abcam, USA), and Donkey anti-Rabbit DL488 (dilution 1:50, Ab96919, Abcam, USA) were used as secondary conjugated antibodies for visualization. Propidium iodide (dilution 250 µg/ml, ab14083, Abcam, USA) and 4’,6-diamidino-2-phenylindole, dihydrochloride (DAPI) (Thermofisher, USA) was used as nuclear staining. Briefly, slides were incubated with primary antibody overnight at 4°C and protected from the light. Hence, primary antibody was washed away by rinsing thrice in PBS, and samples were incubated with a mixture of secondary antibody and nuclear staining for 30 minutes at RT. Slides were mounted with ProLong™ Glass Antifade Mountant (Thermofisher, USA) and stored at 4°C.

### Quantitative imaging analyses

Images of samples from histochemistry were acquired using an EVOS™ M5000 Imaging System (Thermofisher^TM^, USA), equipped with a 10X Objective, fluorite, LWD, 0.30NA/7.13WD. Brightness, contrast, and exposure were maintained constant across all samples analyzed. Images of samples from immunohistochemistry were acquired using a TCS SP5 confocal microscope (Leica, Germany) equipped with a Leica HCX PL Apo 40x/1.3 Oil objective (Leica, Germany). Sequential scanning was performed adopting a resonant scanner set at 8000 MHz frequency. Both the transmission and the fluorescence main intensities were quantified on 8 different images using the following iterative process developed in Fiji software v2.14 on each image:

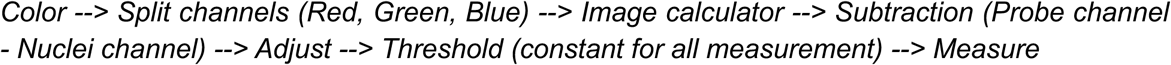

### Osteoarthritis scoring

The tissue morphology of each sample was scored according to OARSI guidelines for the OA-like induced structural modifications. The procedure was based on unbiased observations performed by 5 different scientists on sliced samples stained with SafO-FG, following a two-step approach according to OARSI guidelines^63^. Briefly, an OA grade (0 to 6) was preliminarily assigned to each sample (D0, -L-R, +L-R, +L+R) based on visual modifications occurring both on tissues (e.g., fissures, fibrillation), and cells (e.g., ghost lacunae, chondrocyte hypertrophy). Then, a stage (0 to 4) was assigned to each sample depending on the horizontal extension of damage irrespective of the OA grade. Finally, the score was calculated using the following equation:

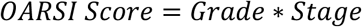

The procedure was performed following a single-blinded approach.

### Statistical analysis

Statistical analysis was run in GraphPad Prism 10.0 performing preliminary tests on data distribution which were deemed normal or non-normal. All comparisons were conducted running either a one-way ANOVA followed by Friedman’s *post hoc* test (non-parametric), or a two-way ANOVA followed by Tukey’s multiple comparisons *post hoc* test (parametric). All variables are presented as mean ± standard deviation (SD), obtained from different replicates of tissues from three different donors. Differences are considered significant with a 95% confidence interval (CI) (p < 0.05).

Sample size (n = 3 biological donors with technical replicates) was determined based on established *ex vivo* cartilage studies^64^ showing sufficient power to detect 30% differences in matrix composition with 80% power at α = 0.05. This approach balances the biological variability across donors with the technical precision of multiple plugs per donor, according to the 3R (refine, reduce, reuse) principle.

## III. Results

### LPS induces reversible OA-like modifications in our model

LPS stimulation impactfully induced an OA-like articular fibrillation in the superficial zone of our model (**Fig. 2 E, F, O, P**), which was completely reversed by Rapamycin (**Fig. 2 G, H, Q, R**). Alcian Blue staining showed reduced intensity in the superficial layer of LPS treated samples (+L-R) (**Fig. 2 E, F**), suggesting GAG loss (**Fig. 2 E, F**, section above dashed red line). Differently, when LPS treated explant were subsequently treated with Rapamycin (+L+R) (**Fig. 2 G, H**) they exhibited an intense blue staining suggesting GAG recovery post-Rapamycin treatment compared to the +L-R (**Fig. 2 E, F**) condition that did not receive any Rapamycin and more similar to the -L-R (**Fig. 2 C, D**) and D0 (**Fig. 2 A, B**) controls. The complete GAG content recovery visible in +L+R compared to -L-R points to an effect of the Rapamycin treatment. Moreover, the minimal disruption of GAG content in D0 and -L-R explants confirms non-invasiveness and robustness of our *ex vivo* model. Analogously, H&E staining highlights an overall loss of native morphology with disrupted chondrocytes and decellularized cartilaginous lacunae in the superficial zone of +L-R (**Fig. 2 O, P, black, and** red asterisks respectively) compared to -L-R (**Fig. 2 M, N**) and D0 samples (**Fig. 2 I, L**). However, a complete recovery from fibrillation-like lesions is noticeable in +L+R samples treated at day 7 (**Fig. 2 Q, R**). Here, the superficial chondrocytes rearrange linearly following the articular rim, similarly to the healthy anatomy of articular cartilage.

**Figure 2:**
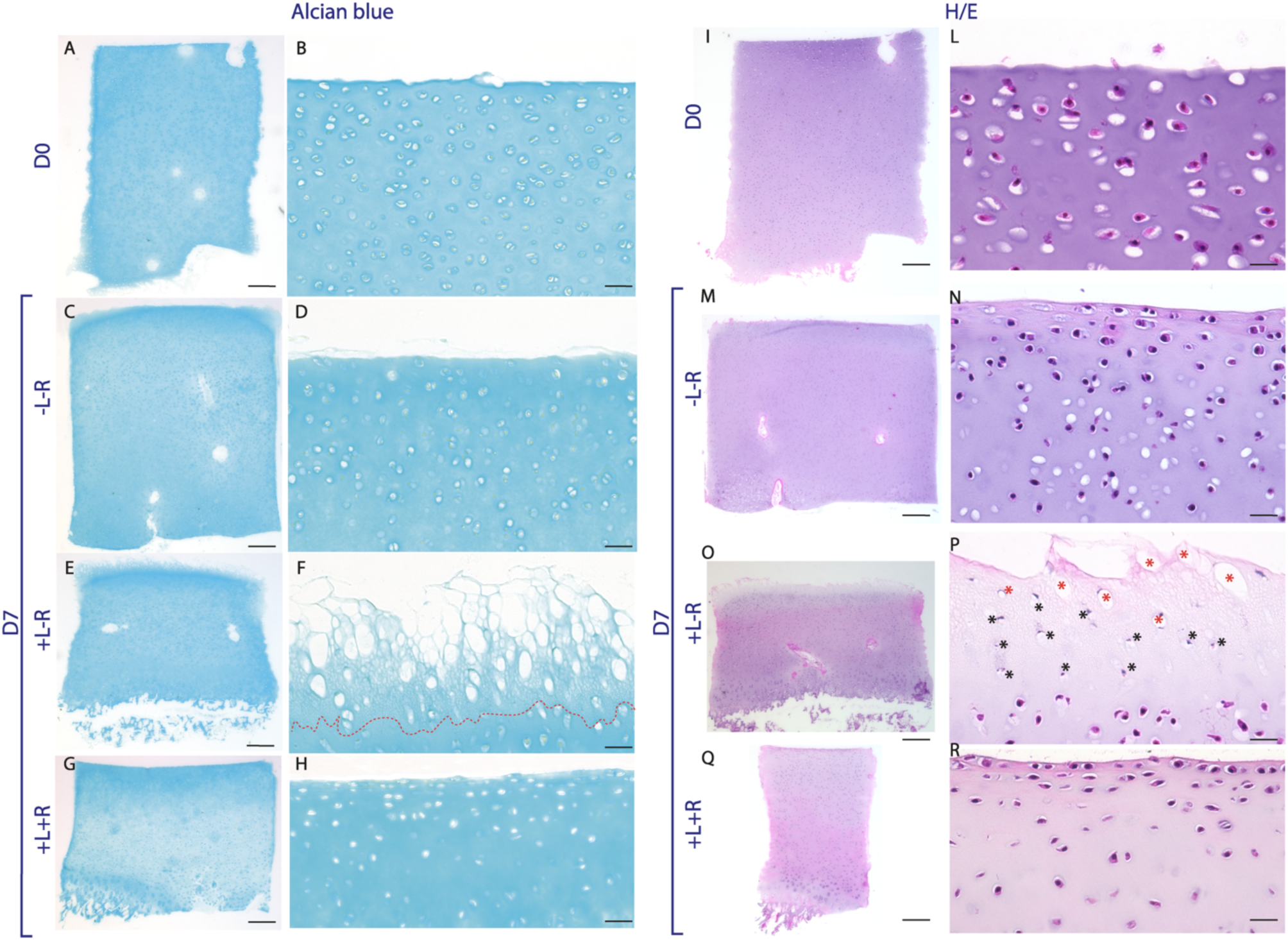
Alcian blue (A-H), and hematoxylin/eosin (I-R) staining showing OA-like changes in our model. There is reduced GAG content in the superficial zone (E, F, zone above the red dotted line) post LPS stimulation. Disrupted chondrocytes (black asterisks) and decellularized lacunæ (red asterisks) are noticeable in LPS-treated samples (O, P). Scale bar = 200 µm (left columns), and 20 µm (right columns).

### Rapamycin restores LPS-induced type II collagen loss

Results from quantitative immunohistochemistry show a substantive presence of type II collagen correlating with healthy hyaline cartilage in untreated samples (-L-R) (**Fig. 3 A, D**). LPS stimulation (+L-R), however, significantly reduces type II collagen in the ECM (**Fig. 3 B, E, G)**. Strikingly, Rapamycin treatment (+L+R) reverses this trend, maintaining high levels of type II collagen in the ECM (**Fig. 3 C, F**).

**Figure 3:**
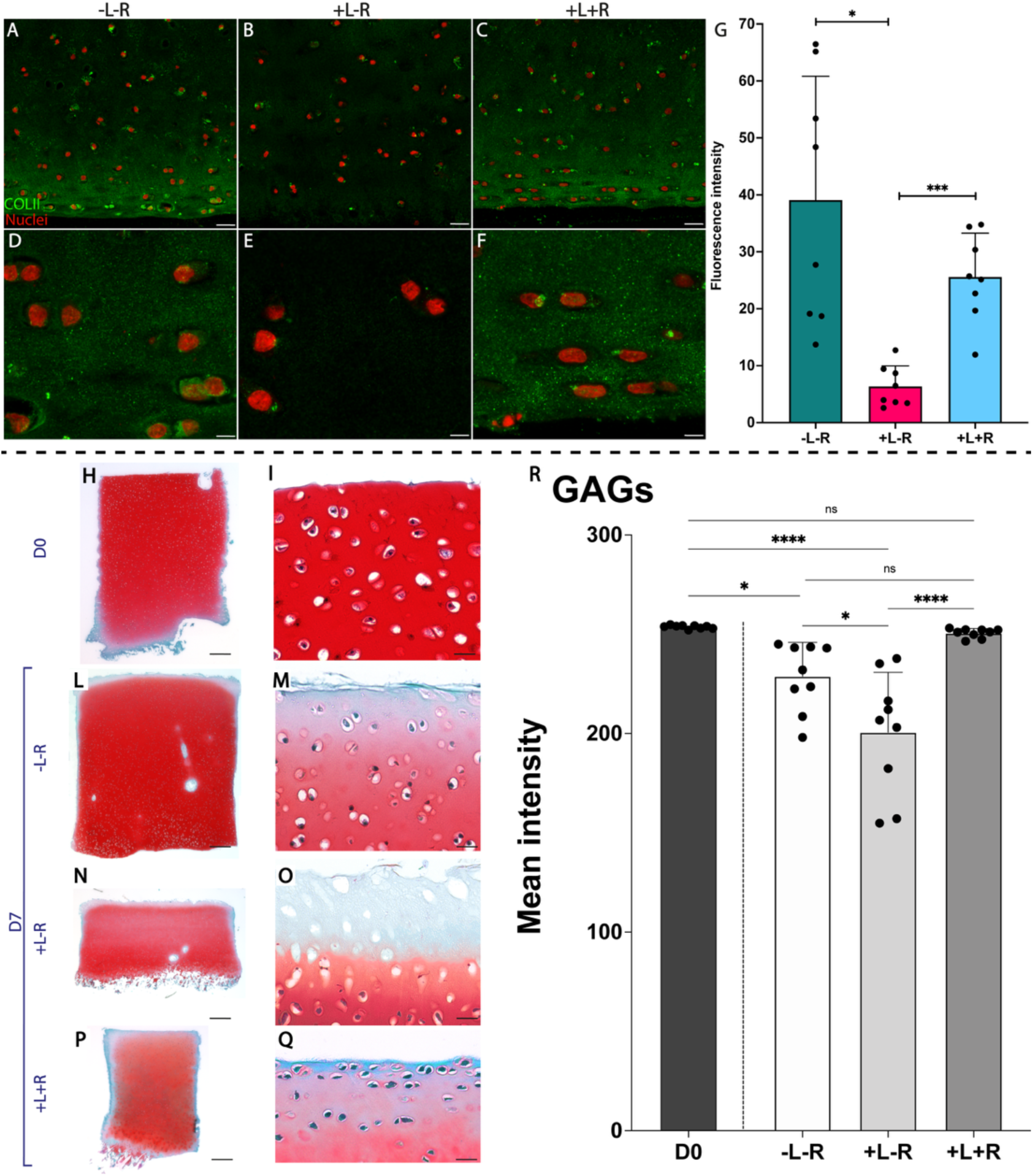
Upper panel - Confocal images of type II collagen visualized with a green labeled secondary antibody on histological sections. Collagen II is reduced in LPS stimulated samples (+L-R) (B, E, G). The treatment with Rapamycin (+L+R) (C, F) promotes an increase of collagen II production, which is visible at level comparable to untreated samples (-L-R) (A, D, G). One-way ANOVA followed by Tukey’s post hoc test, n = 8, * = p < 0.05, ** = p < 0.01, CI = 95%. Lower panel - SafO-FG staining of cartilage samples at baseline (D0) (A,B), and after 7 days of culture without LPS and Rapamycin (-L-R) (C,D), with LPS but without Rapamycin (+L-R) (E, F), and with both LPS and Rapamycin (+L+R) *(G-H). The red staining intensity quantitatively correlates with GAGs content. Scale bar = 200 µm (left columns), and 20 µm (right columns). GAGs quantitative analysis performed on SafO-FG stained samples reported* as mean red staining intensity *(I),* two-way ANOVA followed by Tukey’s *post hoc* test, n = 9, * = p < 0.05, **** = p < 0.001, ns = not significant, CI = 95%.

### OA-like modifications are quantifiable using OARSI scoring

Our LPS-mediated stimulation protocol successfully induced macroscopic modifications on the plugs, similar to OA cartilage fibrillated morphology *in vivo*. Overall, results from SafO-FG staining (**Fig. 3, lower panel**) are consistent with the findings obtained through Alcian Blue and H&E (**Fig. 2**). Precisely, there is noticeable and significant GAG loss in +L-R (**Fig. 3N, O, R**) compared to -L-R (**Fig. 3L, M, R**) and D0 (**Fig. 3H, I, R**). Furthermore, Rapamycin treatment induces recovery of LPS-dependent GAG loss (**Fig. 3P, Q**). LPS-treated samples do show significant GAG loss compared to the native D0 (**Fig. 3R**). Notably, the -L-R sample (**Fig. 3L, M**) remains undamaged overall, appearing morphologically comparable to the D0 native tissue, further confirming the robustness of our *ex vivo* model (**Fig. 3H, I**). Most importantly, the SafO-FG staining highlighted well-defined morphological parameters in our model, making it possible to perform scoring following the standardized protocol described in the OARSI guidelines^63^.

After independent assessment from 5 different observers (**Fig. 4 A**), the +L-R samples show a significantly higher score, suggestive of cartilage degradation, compared to both +L+R and D0 samples (**Fig. 4 D**). Remarkably, +L-R leads to significantly higher grade (**Fig. 4 B**) and stage (**Fig. 4 C**) values compared to D0 samples using the OARSI evaluation process. There was no significant difference between -L-R and D0 samples in grade, stage, or score, further pointing to the robustness of the explant culture. Thus, these findings demonstrate an induction of OA-like modifications in our *ex vivo* model, which is fully reversible after Rapamycin treatment.

**Figure 4:**
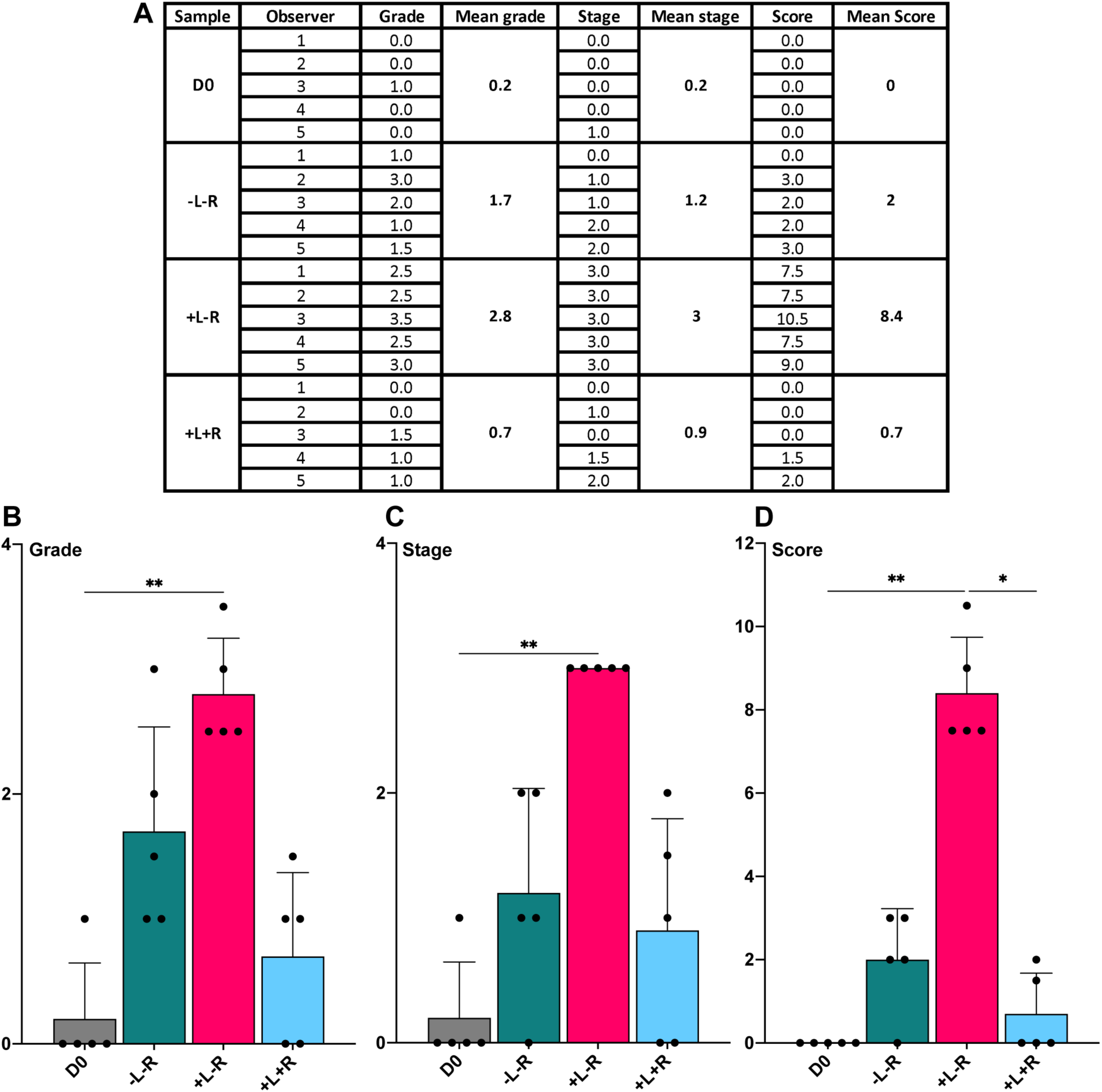
*Grading, staging and scoring of Osteoarthritis performed by 5 different observers following the methodology described in the OARSI guidelines (A). Statistical analyses showing significant differences of grade (B), stage (C), and score (D) between different experimental conditions,* one-way ANOVA followed by Friedman’s *post hoc* test, n=5, * = p < 0.05, ** = p < 0.01, CI = 95%.

### LPS stimulation induces the ROMO1 pathway activation

LPS stimulation induced ROMO1 activation in +L-R samples (**Fig. 5 B, E**), while both +L+R, and -L-R samples (**Fig. 5 C, F, and A, D respectively**) showed significantly lower levels of ROMO1. Remarkably, the ROMO1 levels in +L+R samples decreased to values comparable to the constitutive levels seen in untreated -L-R samples (**Fig. 5 G**).

**Figure 5:**
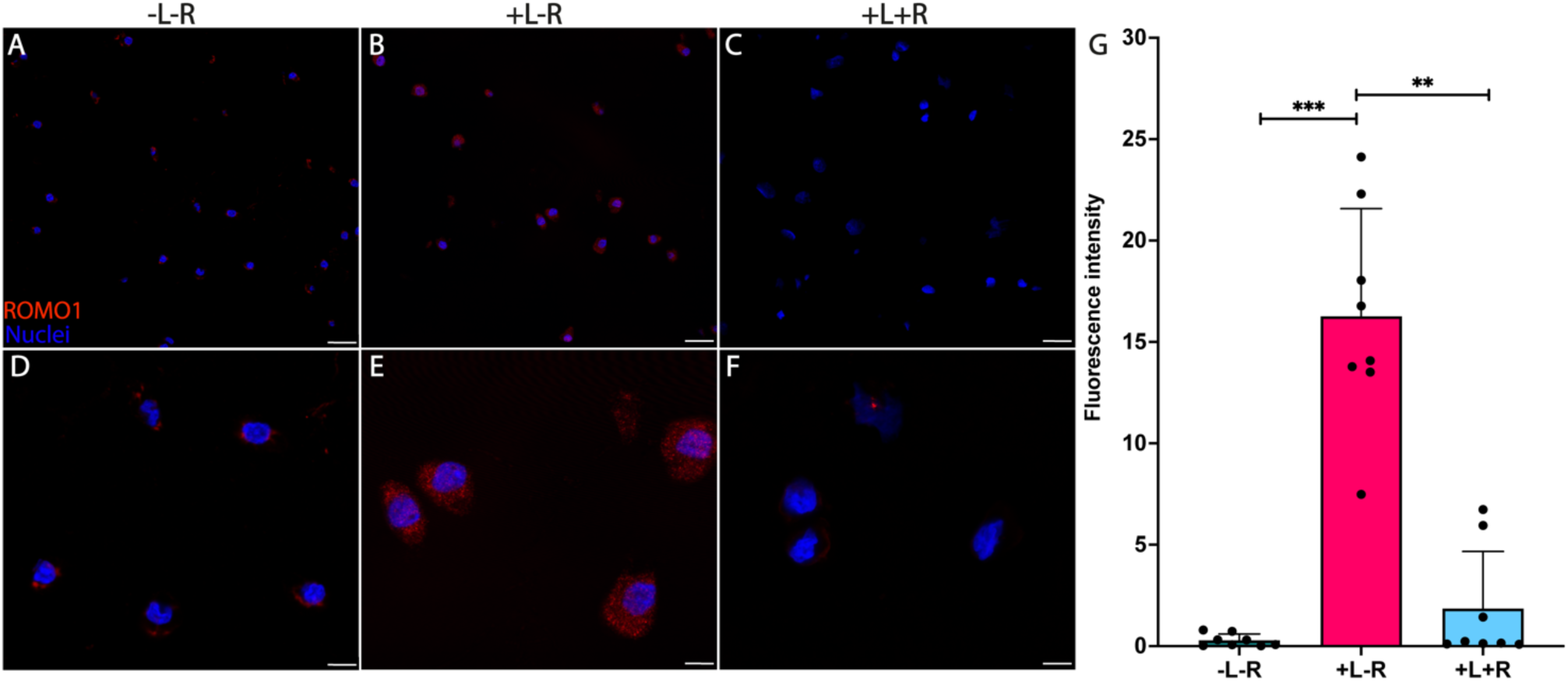
Confocal images of ROMO1 protein complex visualized with a secondary antibody (red) on histological sections. ROMO1 expression is increased in LPS stimulated samples (+L-R) (B, E). The treatment with Rapamycin (+L+R) (C, F) promotes a decrease of the ROMO1 activation, which is higher compared to untreated sample (-L-R) (A, D). Scale bar = 20 µm (A, B, C), 10 µm (D, E, F). Quantitative measurement of ROMO1 fluorescence intensity (G), Friedman’s test followed by Dunn’s *post hoc* test, n = 8 * = p < 0.05, ** = p < 0.01, CI = 95%.

### Rapamycin treatment on *ex vivo* LPS-treated cartilage plugs displays modulation of autophagy-related pathways

The stimulation with LPS interacts with autophagy-related pathways, by modifying the production of key factors such as LC3b and ULK1. The detection via immunohistochemistry shows that both LC3b and ULK1 are physiologically present at steady-state in untreated tissues (-L-R, **Fig. 6 A, D, 8 H, M**). The LPS stimulation induces a significant reduction of LC3b and ULK1 in stimulated cartilage, +L-R, which is clearly indicated by the reduced green fluorescence (**Fig. 6 B, E, 6 I, N**). Interestingly, Rapamycin treatment induces a strong reversal of this phenomenon in LPS-treated samples (+L+R), leading to significantly higher LC3b (**Fig. 6F**) and ULK1 (**Fig. 8O**) expression compared to the LPS-treated samples (+L-R). Moreover, in Rapamycin treated samples, the LC3b signal is also significantly higher compared to time-matched untreated controls (-L-R) (**Fig. 6G**), while the ULK1 signal is not higher (**Fig. 8Q**).

**Figure 6:**
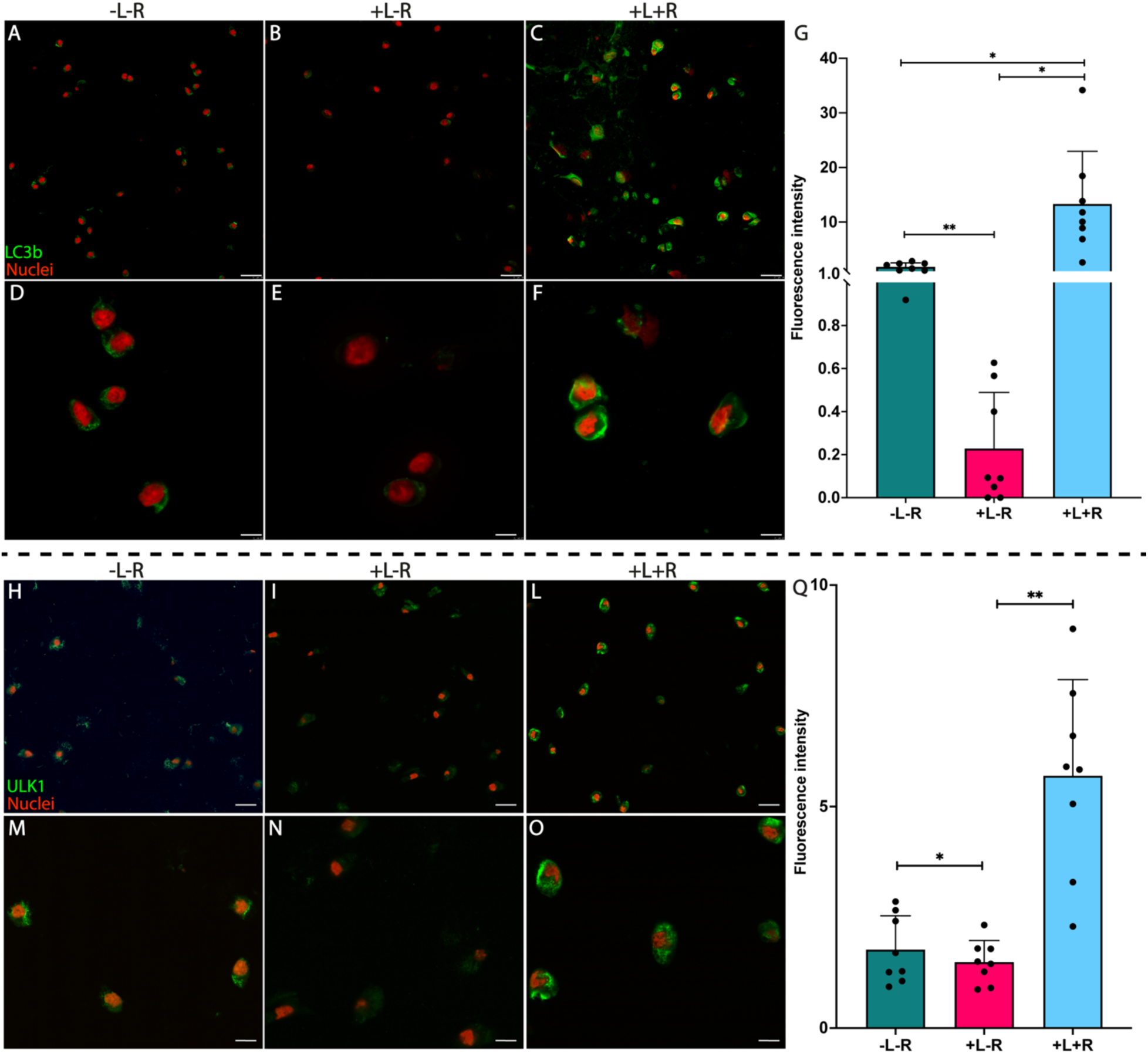
*Upper panel -* Confocal images of LC3b protein visualized with a secondary antibody (green) on histological sections. LC3b expression is reduced in LPS stimulated samples (+L-R) (B, E) but then recovered with Rapamycin treatment (+L+R) (C, F). Rapamycin treatment(+L+R) (C, F) also leads to increased LC3b expression compared to untreated samples (-L-R) (A, D). Scale bar = 20 µm (A, B, C), 10 µm (D, E, F). Quantitative measurement of LC3b fluorescence intensity (G), one-way ANOVA followed by Tukey’s post hoc test, n = 8, * = p < 0.05, ** = p < 0.01, CI = 95%. Lower panel - Confocal images of ULK1 protein visualized with a secondary antibody (green) on histological sections. ULK1 expression is reduced in LPS stimulated samples (+L-R) (B, E). Rapamycin treatment (+L+R) (C, F) promotes higher ULK1 expression compared to LPS-stimulated (+L-R) (B, E) and untreated samples (-L-R) (A, D). Scale bar = 20 µm (A, B, C), 10 µm (D, E, F). Quantitative measurement of ULK1 fluorescence intensity (G), one-way ANOVA followed by Tukey’s *post hoc* test, n = 8, * = p < 0.05, ** = p < 0.01, CI = 95%.

## IV. Discussion

This study aims to study the downstream inflammatory and matrix changes in cartilage due to TLR4 signaling and compare the morphological changes in cartilage post LPS insult as well as in recovery, after treatment with an anti-inflammatory drug, Rapamycin. We develop a realistic stimuli-responsive *ex vivo* OA model that could also be used to study the gut-joint axis. The model is based on native cartilage explants, in this specific case porcine as proof of concept, and is conceived to mimic the pathophysiology of OA, acting as a bench test for new therapeutic strategies.

By using LPS as the single inflammatory-trigger to induce an OA-like state via TLR-4 pathway activation^11^ in an *ex vivo* culture model, our goal was to overcome limitations of routinely adopted protocols based on stimulation with pro-inflammatory cytokines (i.e., IL-6, TNF-a)^61^, which activate positive-feedback loops promoting further release of the same cytokines^62^, making it impossible to distinguish exogenous from endogenous drivers of inflammation. Besides presenting a realistic stimuli-responsive *ex vivo* model based on LPS stimulation to induce OA-like modifications, this study also assessed the effect of subsequent treatment with Rapamycin leading to phenotype recovery. We successfully recapitulated several key aspects of OA pathophysiology *in vivo*, including extracellular matrix degradation, autophagy suppression, and oxidative stress. Our results demonstrate that LPS stimulation effectively causes GAG and collagen loss in ECM, leading to an increase of ROMO1 (correlated with ROS and oxidative stress), and a decrease of autophagy-related markers (ULK1, LC3b), which are all reversed via Rapamycin treatment.

Our LPS-treated plugs showed decreased GAG and collagen content compared to the control plugs. These results align with literature where inflammatory stimulation leads to the degradation of the ECM and increase in matrix remodeling proteins^10^. GAGs are among the most abundant ECM components in cartilage and reduced GAG production in chondrocytes^12,65–67^ is one of the most common biochemical signs of OA. Noteworthy, GAG reduction is reported to be more evident in the superficial zone, which *in vivo* is the first cartilage region to sustain damage during OA ^68^. Furthermore the morphological analysis of LPS-treated plugs shows superficial lacunae decellularization and disrupted chondrocytes in the inferior adjacent area, which in *in vivo* models have been attributed to a chondrocytes attempt to repair the damaged smooth cartilage surface^69^. The morphology of our model is strikingly comparable to those described from Pritzker and colleagues^63^, where the OARSI scoring system was also adopted. Furthermore, collagen degradation is another notable characteristic of OA cartilage^70,71^ and our LPS-treated plugs recapitulates it well, in agreement with the *in vitro* literature on the use of LPS to stimulate osteoarthritic conditions^72^.

Notably, the model demonstrates responsiveness to Rapamycin treatment, a macrolide with well-established beneficial effects toward OA^73,74^. We were impressed to notice how the OA-like conditions we induced in our model were entirely reversed with Rapamycin treatment. In fact, as a result of the treatment with Rapamycin treatment the lost ECM was fully recovered both in terms of GAGs content and gross morphology, as evidenced by our histologies. It is worth pointing out that time-matched untreated samples at day 7 are comparable to native tissues at day 0, suggesting that our culture protocol does not negatively affect native GAGs content or macroscopic morphology after 7 days. Interestingly, we observed that a 3-day window of treatment is sufficient for recovery, which is consistent with the evidence described by Ikenoue *et al*^75^ who reported a fast increase of type II collagen and aggrecan from human chondrocytes after being exposed to pro-regenerative stimuli. Taken together, these observations further validate our *ex vivo* model as a suitable approach to test signaling pathways in OA and potential of therapeutics.

As expected, our results show that LPS stimulation led to ROS production as evidenced by the upregulation of the ROMO1 protein complex. Upregulation of ROS is a key element of OA as it can lead to cartilage degradation and activate the PIK3/AKT/mTOR pathways^44–48^. Rapamycin treatment was able to reverse ROS production via downregulation of ROMO1. This phenomenon has been previously observed in literature *in vivo* in aged mice^76^ as well as using a surgically induced OA mouse model where Rapamycin led to a decrease in IL-1β and ADAMTS-5 expression, both of which are proteins known to stimulate ROS production^73^.

LPS stimulation also led to a clear decrease in autophagy in the *ex vivo* porcine plugs. This was evidence via decreased ULK1 and LC3b expression post stimulation, both of which are expressed in healthy murine articular cartilage. These *ex vivo* results agree with Carames *et al* which showed that surgically induced OA in mouse cartilage leads to decreased ULK1 and LC3 expression^38^. The phenomenon of decreasing autophagy in OA conditions suggests a defect in the cellular remodeling process, leading to the accumulation of damaged macromolecules and an increased likelihood of aging-related diseases^38,77^. Thus, our *ex vivo* model, even with just a short LPS inflammatory stimulation, was sufficient to inhibit autophagy, mimicking *in vitro* the outcomes observed in mouse models. Furthermore, Rapamycin administration resulted in the recovery of ULK1 and LC3 expression in LPS-treated plugs. Similar to the results in our *ex vivo* model, Tang *et al* showed that LPS-treated human primary chondrocytes exhibited decreased autophagy and increased ECM degradation^32^. These authors used diindolylmethane, an inhibitor of inflammation and apoptosis, in their 2D cell culture model which leads to decreased extracellular matrix degradation and upregulation of autophagic signals, paralleling our results using Rapamycin. Moreover, they also observed an increase in LC3 II or LC3b expression post inhibitor use suggesting that LPS-induced chondrocyte autophagy and ECM degradation can be regulated via the PI3K/AKT/mTOR- autophagy axis.

This work focused on creating a realistic *ex vivo* OA model that mimics the pathophysiology of OA, allowing for mechanistic hypotheses investigations, and acting as a bench test for new therapeutic targets. Our model -successfully recapitulated some key features of OA, including ECM degeneration at the microscale in porcine cartilage plugs. Moreover, we demonstrated that the model behavior is driven by well-known OA-related molecular pathways, including TLR4 activation, mTOR signaling, and autophagy. Although our model may be further improved, we believe that it has the potential to pave the way to new studies focused on the pathophysiology of OA, and the ability to test therapeutic approaches *ex vivo*. Advantageously, our model offers rapid readouts (7 days) compared to surgically-induced models *in vivo* (8 – 12 weeks), while offering a realistic 3D environment and avoiding confounding variables, such as undesired inflammatory responses triggered by altered biomechanics.

However, while our model successfully replicates key OA characteristics, several opportunities exist to expand its utility in the near future. First, the inclusion of TLR4-specific inhibitors (e.g., TAK-242) would ultimately confirm the pathway specificity, distinguishing TLR4-mediated from unspecific LPS effects. In this study, we intentionally focused on the downstream autophagy markers (ULK1, LC3b) and functional matrix outcomes rather than on direct mTOR phosphorylation, as we believe that these better reproduce the biological responses to cartilage homeostasis. However, future refining studies could take advantage from phospho-mTOR immunohistochemistry to provide a complementary pathway validation. Second, implementing analyses capable of assessing chondrocyte proliferation would better quantify the regenerative response to Rapamycin treatment. Although the current model focuses on matrix recovery and autophagy restoration, the study of chondrocytes’ proliferation kinetics would give insight on the optimal dosing-windows for an efficient and safe clinical translation. Finally, extending the protocol to human OA cartilage explants obtained from patients undergoing arthroplasty could help confirm that OA shows sensitivity to mTOR inhibition in humans. Such data might open the way to clinical trials exploring mTOR-targeting therapies and to the development of OARSI-validated protocols for personalized *ex vivo* drug screening on patient-derived tissue. Together, these approaches help promote the development of new methodologies based on precision medicine to ultimately accelerate the clinical translation of new drugs candidates for OA.

## Acknowledgments

This work was partially supported by the OActive project, funded by the EU Horizon 2020 Programme under grant agreement No. 777159 (funding scheme: H2020-SC1-PM-17-2017). Additional funding was provided by the ON Foundation (Lucerna, CH) (Starting Grant No. 22-006). This study was also supported in part by the National Institute of Dental & Craniofacial Research (NIDCR) of the National Institutes of Health (NIH) under Award Number T90DE030854 (or R90DE031532) and the Center for Innovation & Precision Dentistry (CiPD) at the University of Pennsylvania and NIH F32 DE035372.

