## Supplementary figures and images for "A stimuli-responsive *ex vivo* model of osteoarthritis demonstrates TLR4-mediated cartilage degradation and a Rapamycin-induced fast matrix recovery"

### Supplemental figure 1

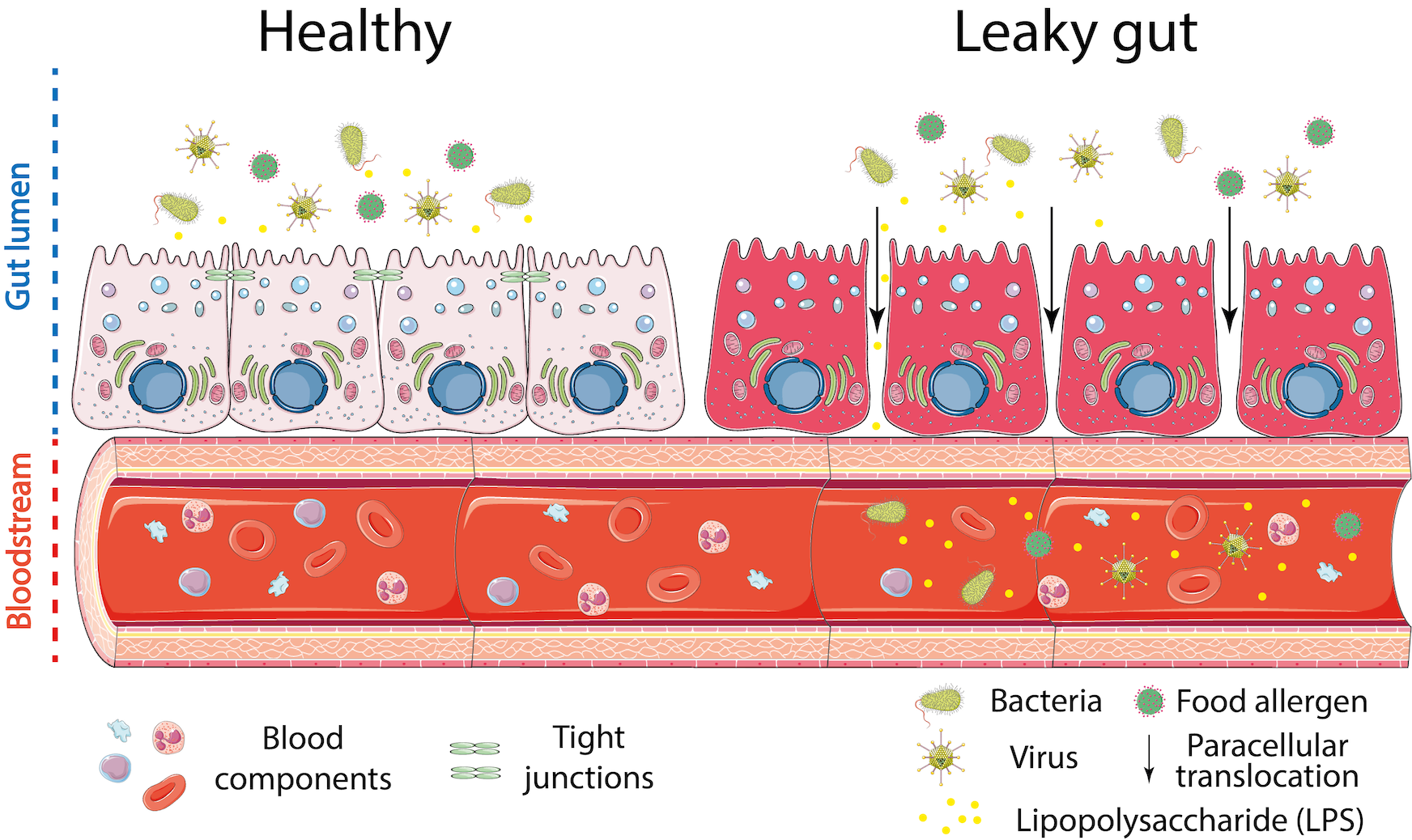

### Supplemental figure 2

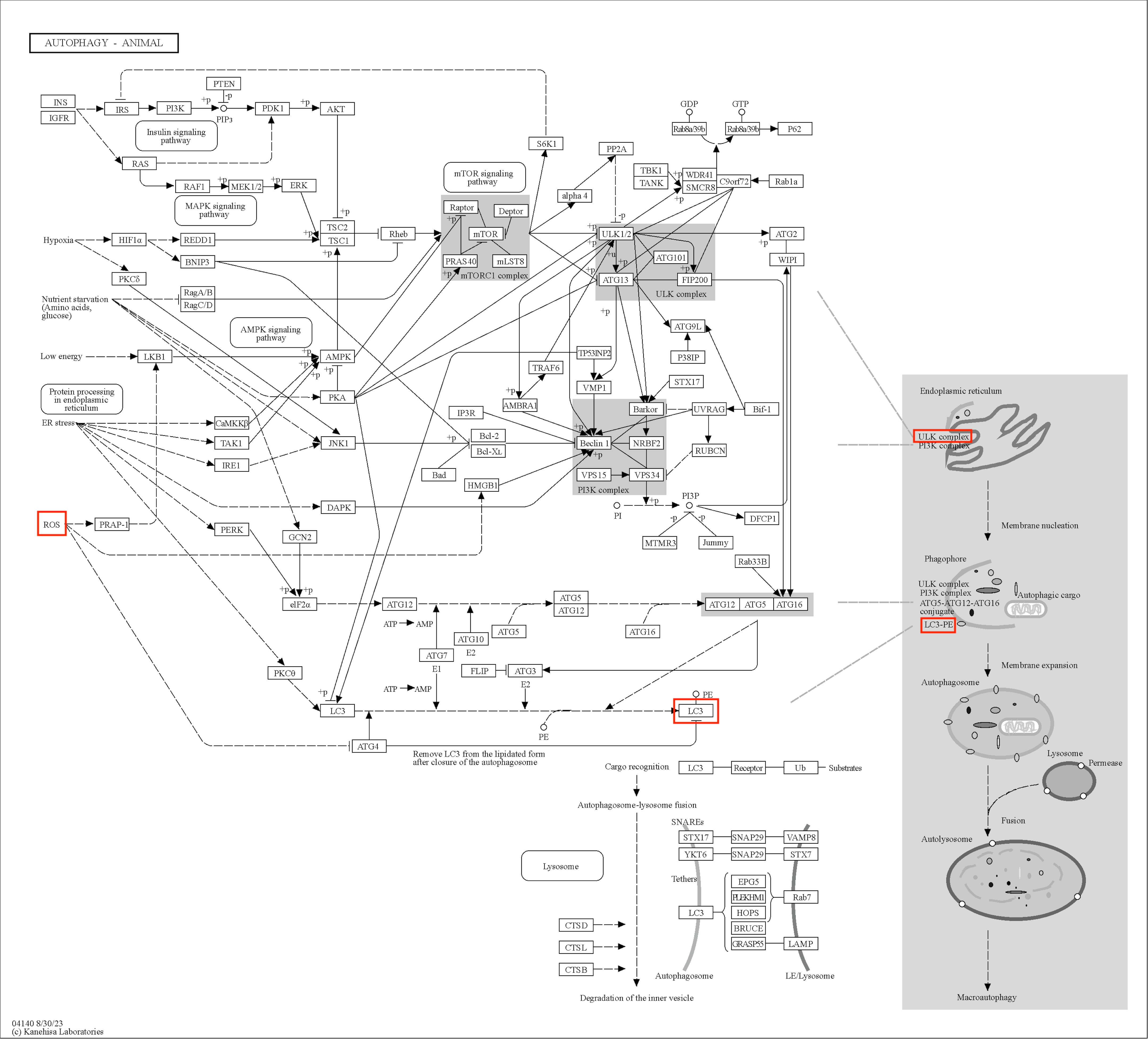

### Supplemental figure 3

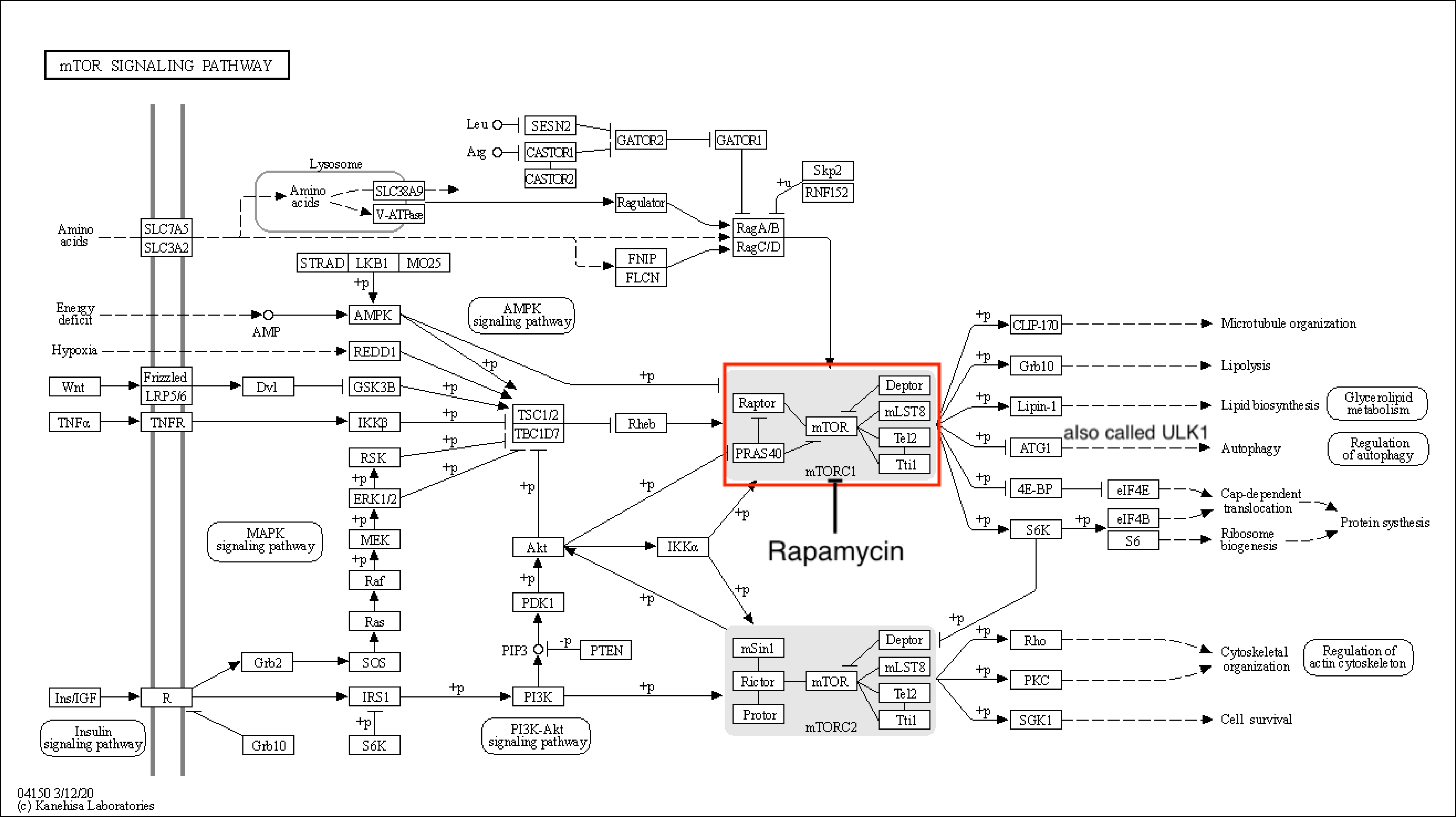

### Supplemental figure 4

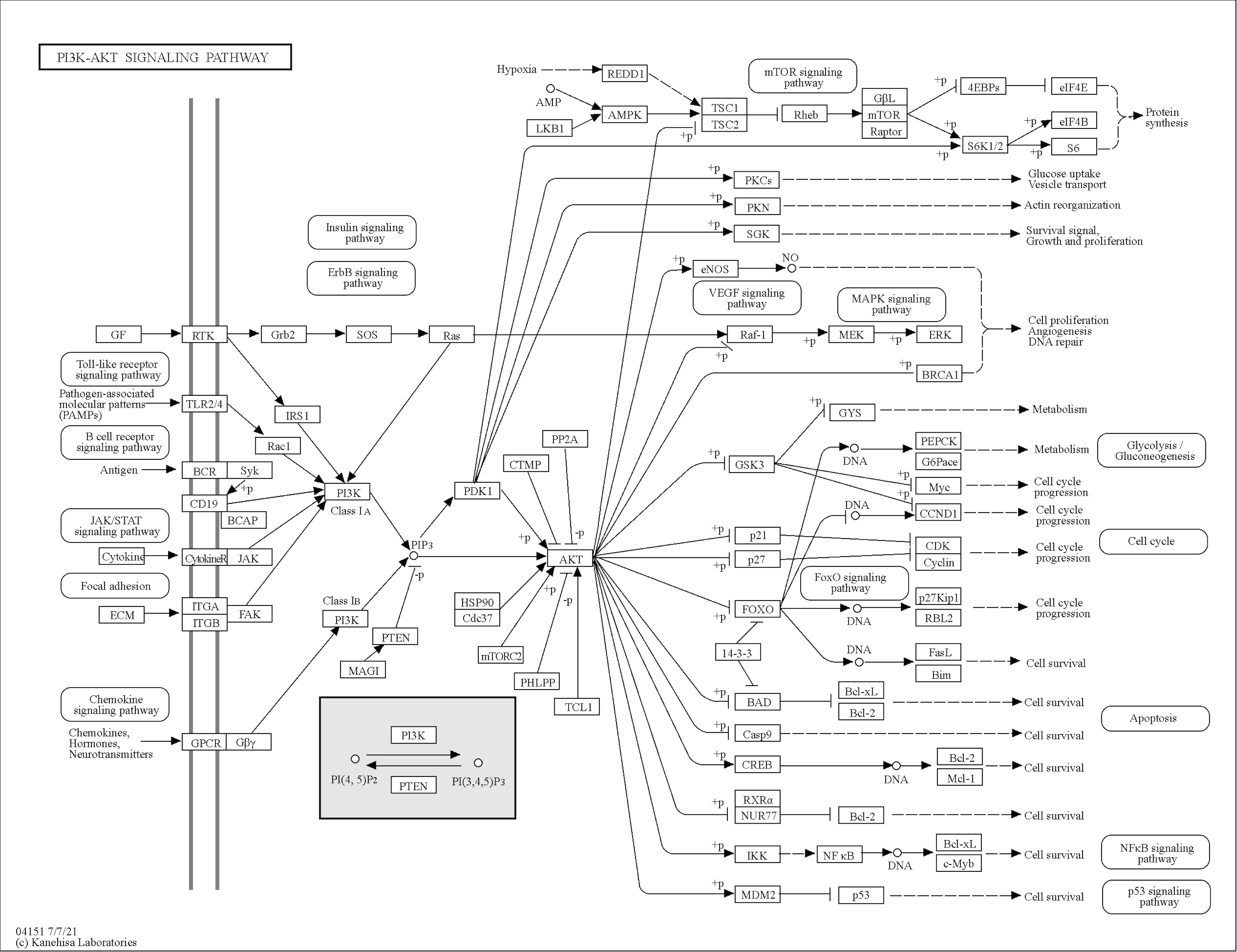
